# Production of Hybrid Rice seeds using environment sensitive genic male sterile (EGMS) and basmati rice lines in Kenya

**DOI:** 10.1101/755306

**Authors:** Njiruh Paul Nthakanio, Kariuki Simon Njau

## Abstract

Photoperiod-sensitive genic male sterile rice (PGMS) lines IR-73827-23-76-15-7 S, IR-75589-31-27-8-33S referred to as P1 and P2, and IR-77271-42-25-4-36S, thermo-sensitive genic male sterile (TGMS) line referred to as T were obtained from International Rice research Institute. These lines, collectively known as environment genic male sterile lines, were sown under greenhouse growth conditions where temperatures were more than 34°C with an objective of inducing complete male gamete sterility in them. Results indicated that high temperature growth conditions induces complete male gamete sterility in both the PGMS and TGMS lines. The impact of this is that, it will be possible to produce pure basmati hybrid rice seed in the tropical regions without contamination with pure breed lines. The male sterile PGMS/TGMS were pollinated with pollen from basmati370 and 217 grown under natural conditions and some hybrid seeds were obtained. This shows that high temperature emasculated the male gametes but not female ones. The conclusion is that it is possible to induce complete male gamete sterility in PGMS and TGMS under greenhouse in tropical growth conditions, and to produce hybrid rice seeds. This makes basmati hybrid rice seed production in Kenya a viable venture.

## Introduction

The World rice production was about 503.6million tons in 2017 (1). This is below consumption that was 505.8 million tons in the same period. In Kenya, rice consumption is over 580,000 tonnes against a total production of about 149,000 tonnes (2). The deficit, which is valued at over Kenya shillings Seven billion is imported (2). Basmati rice is preferred by many consumers compared to non-aroma varieties because of its good cooking traits (3). However, in Kenya, basmati yields only 3.6 to 4.0tones per hectare (4). This is quite low and it has contributed to keeping its prices high. Over the years, rice breeding has gone through a breeding paradigms with emphasis of high yield (HYV) semi-dwarf varieties (5). The major shift came with the green revolution which brought about IR8 variety in 1966 with the dwarf gene *sd-1* (6) which raised the yield to over 6 tones per hectare (7).

Hybrid rice technology was introduced in 1970s (8, 9) to improve yield above dwarf lines. Heterosis improved yields in rice (10, and hybrids lines are reported to have a 20-25 percent yield advantage over pure breeds (11). However, some advantages of this have been eroded by diseases such as blast (12). To overcome this, green super hybrid technology has been adopted that further increased realizable rice yield per hectare by 12% above the normal hybrids (13). Advances in green super hybrid technology started in China in 1996 and it targeted raising rice grain yield from about 10 tones to about 17 tones per hectare (14). The yield was realizable by combining hybrid vigour and good agronomic traits such as disease resistance (15). According to Yuan Longping (16), rice yield in China stands at about 17 tones per hectare.

A number of approaches have been used in hybrid rice production that include the three line system, which utilizes cytoplasmic male sterility (CMS) (17) and the two line hybrid system that are referred to as environment sensitive genic male sterility (EGMS) (18–20). Among the EGMS is the photoperiod-sensitive genic male sterile (PGMS) rice line that is completely sterile when under 14hours daylight length growth conditions. It reverts to fertility in varying degree when grown under less than 14 hours daylight length conditions (18, 19). Other EGMS are thermosensitive genic male sterile (TGMS) rice lines that is sterile when grown under high temperature and revert to some fertility when grown under low temperature growth conditions (17). In their sterile phase the EGMS rice lines are crossed with a male parent to produce F_1_ (hybrid) seeds (21).

Pure basmati rice yield per hectare is low compared to non-aromatic lines (22) and this has kept its prices high. Basmati370 and 217 varieties are the two major aromatic rice varieties grown in Kenya. Exploitation of hybrid technology to improve their yield is limited (23). Elsewhere, attempts to produce hybrid basmati rice lines have shown a good combining abilities in yield traits (24). In this research object was to produce basmati370 and 217 hybrid rice seeds using two line methods. The yields traits realized were better than that of both purebred lines.

## Materials and methods

### Materials

Environmental genetic male sterility (EGMS) rice varieties, used as female parents were IR-73827-23-76-15-7S (P1) and IR-75589-31-27-8-33S (P2) that are photoperiod sensitive genic male sterile (PGM), and IR-77271-42-5-4-36S (T) that is Thermosensitve genic male sterile TGMS) rice lines which were all imported from IRRI (Philippines) following Kenya Plant Health Inspectorate Service (KEPHIS) importation rules and regulations. The basmat370 and 217, used as male parents, were provided by Mwea Irrigation Agriculture Development (MIAD). Sowing was done at Kenya agricultural and livestock research organization (KALRO) Mwea, which is located at latitude 0.7°S and longitude 37.37E where daylight length and night length are nearly equal (12hour).

### Methods

#### Sowing and testing for adaptability of EGMS lines

Dormancy in EGMS and Basmati rice seeds was broken by submerging them in 2% H_2_O_2_ for 72. A fresh change of H_2_O_2_ was done after every 24 hours. Thereafter, seeds were sown in germinating plates in a nursery until seedlings were 21 days old. Transplanting of seedlings in the field was done at spacing of 20cm x 20cm in growth troughs made of concrete blocks in the greenhouse (GH). Control seedlings were sown outside the greenhouse under natural growth conditions. In each set (inside and outside GH) basmati370 and 217 varieties were sown as the pollen donor parents. The temperature in the GH was maintained to above 35°C and 20°C during the day and night times respectively. During the day temperatures in the green house was regulated downwards by and opening the door or and vents and conserved at night by closing the greenhouse. Plants were allowed to grow till flowering when high temperature treatment was stopped.

#### Screening for male sterility

During the first 10 days after flowering, 10 plants per variety were selected and pollen samples were taken once in every two days for pollen sterility testing. Samples from basmat370 and 217 were used as controls. Three young spikelets (picked from top, middle and bottom) from one panicle per plant were randomly selected from plants in the GH and outside GH. They were fixed in 70% alcohol. Three anthers from each spikelet were stained by placing an anther on a drop of 1% Potassium Iodine (I_2_KI) on a glass slide, then macerated with forceps to release pollen, followed by observations under X10 objective of a light microscope. Pollen fertility was done by counting sterile/abortive (yellow and brown stained) against fertile (dark blue stained) pollen cells. The % fertile pollen (at heading) and fertile spikelets (at maturity) in plants were calculated using the equations below (17);

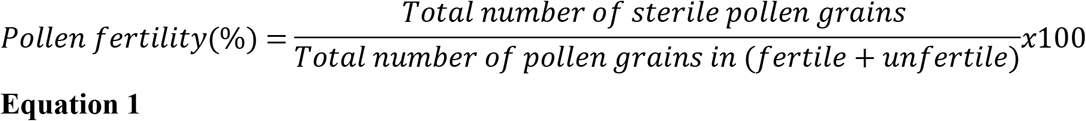

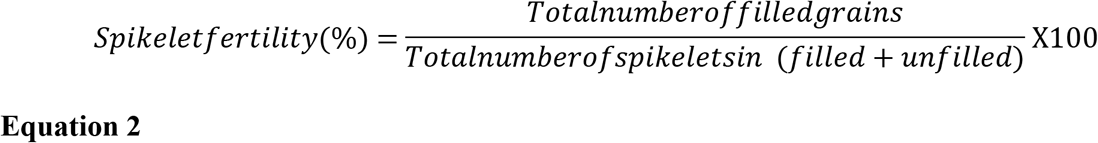

#### Production of hybrid rice

The following crosses were made: P1 X B370, T X B370, P2X B370, P1 X B217, T X B217 and P2 X B217. This was done through crossing EGMS (♀) x Basmati (♂) to obtain F_1_ hybrids. Pollen donors were sown outside the (GH) while female plants (P1, P2 and T) were in GH growth conditions. Sowing was staggered in three stages (from first planting September 23^rd^, 2012) to ensure synchrony during flowering of donor pollen with their recipient. In stage 1 only pollen donors were sown, in stage 2, ten days after stage 1, EGMS (P1, P2 and T) were sown while in stage 3, 20 days after stage 1, only basmati370 and 217 (pollen donors) were sown. At critical sterility point (CRP) which is 30 days before heading, EGMS and basmati parents were exposed to high temperature under GH growth conditions till heading when female plants were pollinated with pollen from male parents. Glumes were clipped at the tips to expose the stigma then pollen from fertile basmat370 and B217 was dusted over the clipped glumes between 11.30pm and 1.30pm. Pollinated panicles were then bagged to prevent unwanted crossings.

#### Evaluation of Agronomic Traits

Hybrids and parental lines were planted in a complete randomised block design (3 blocks with 3 replicates). All standard agronomic practises such as pest and diseases control were done. Yield traits from each sampled plant were measured at the physiological maturity period. These included plant height effective tillers, 1000 seed weight, effective tillers and number of glumes per panicle were determined as described by Virmani, et al. (17). Seeds of each three sampled plants were bulked and a seed counter used to get 1000 seeds that was used to determine grain weight. Days to heading were determined at 50% emergence of panicles starting from the sowing date, while days to maturity was calculated as 30 plus days to 50% heading of each rice line. The percentage seed set rate was determined using equation below;

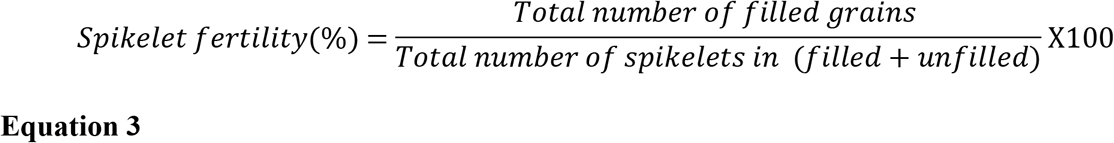

#### Data analysis

Data obtained on temperature, parental pollen viability, height, productive tillers, flowering date, seed setting, panicle length and exertion ANOVA was analysed using SPSS 16.0 statistical package. Numerical data of two environments was expressed in Mean±SD and analysed using students*t*-test for significance. At p≤0.05, mean values, were considered statistically significant.

## RESULTS

### Induction of male sterility in EGMS varieties

The temperatures in the greenhouse (GH) and outside greenhouse (OGH) growth conditions were by average 24°c and 34°c respectively. Within GH growth conditions, line P1, T and P2 recorded pollen fertility of less than 2% while basmati370 and 217 recorded 25% and 21% respectively. All lines grown under OGH conditions recorded over 60% pollen fertility (Fig 1).

**Fig 1:**
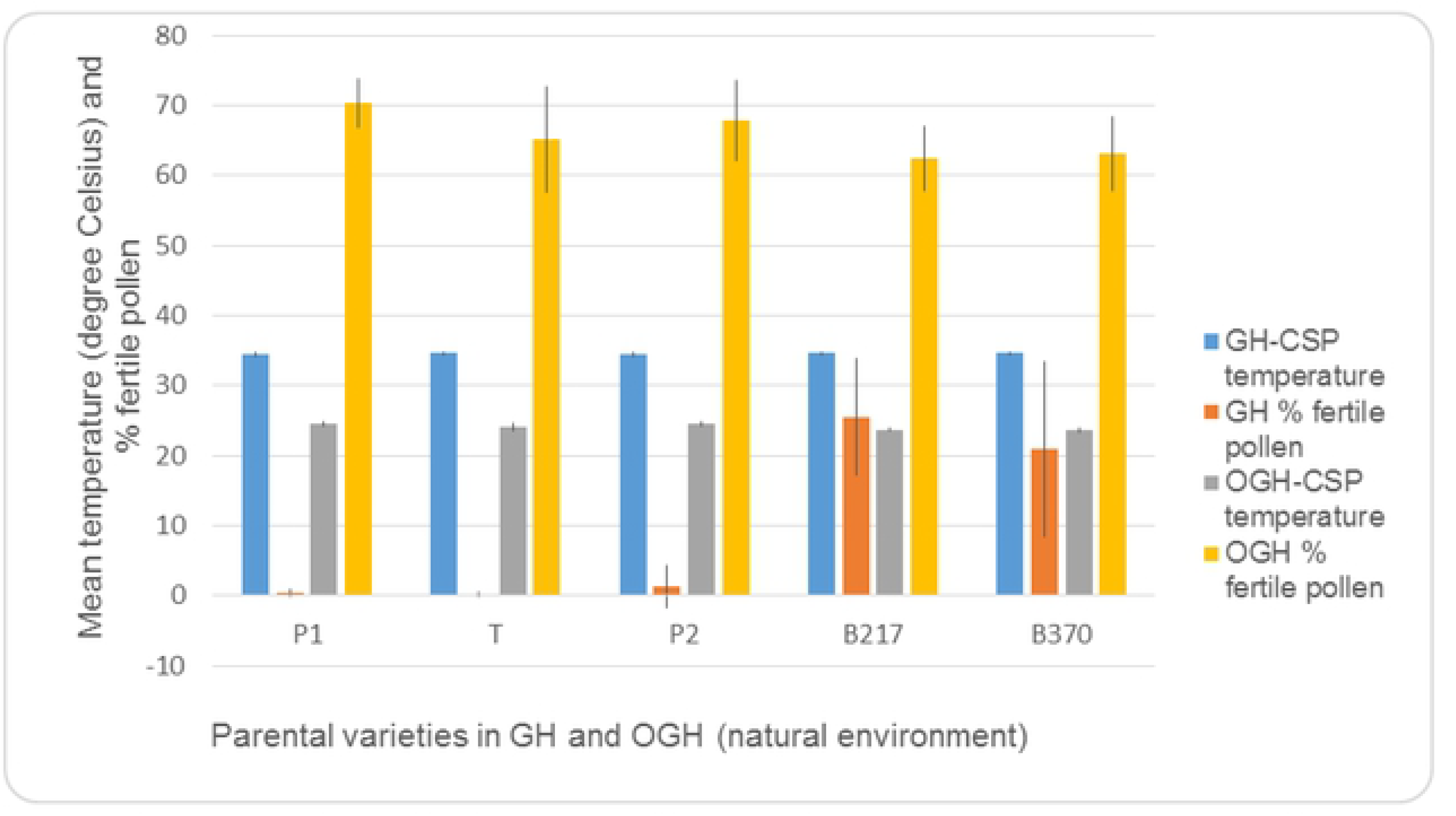
Pollen fertility under GH and OGH growth conditions. Temperature in the green out and outside green house were constant. Scale for temperature is in degrees Celsius and pollen fertility is in %. Lines *P1* and *P2* stand for PGMS, line T stand for TGMS and B stand for *Basmati*.

Most pollen from EGMS grown under GH growth conditions stained yellow or fading blue-black with 1% KI and their anther locules looked empty (Figs 2 a and b). This is what was classified as fertile and abortive pollen respectively (Figs 2 c and d). In the GH environment, all EGMS (P1, P2 and T) pollen was either absent or deformed and stained yellow with 1% KI (Fig 2 c). Some pollen from basmati370 and 217 under GH growth conditions stained blue-black with 1% KI (Figs 2 d).

**Fig 2:**
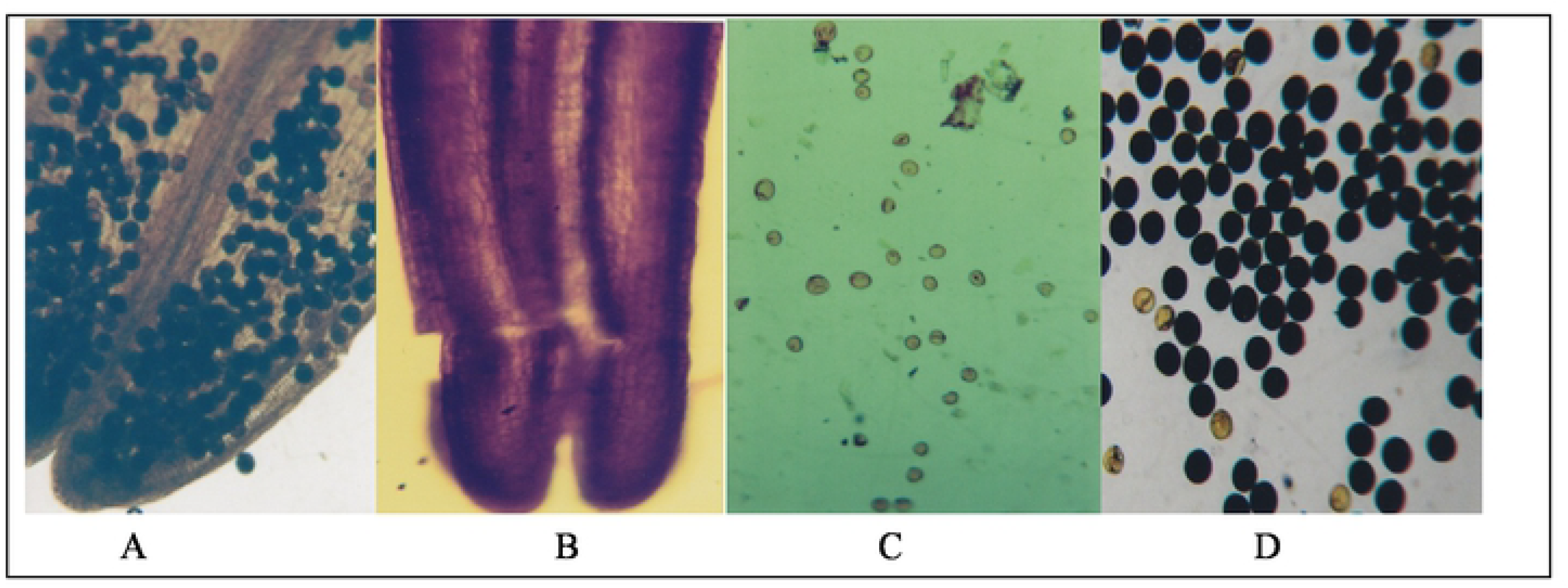
Comparison of pollen fertility (under ×10 magnification) of plants grown in GH and OGH growth conditions. Figs A and B that of glumes take from line B370 and EGMS (P1) under GH growth conditions respectively. Figs C and D are that of P1 and B370grown under GH growth conditions respectively.

The results effectiveness of GH to raise temperature and effectively induce complete sterility in EGMS and subjected to unpaired *T*-test analysis are shown in table 1. Line P1 with 2.4*10^−11^ had the highest pollen sterility rate compared to basmati370 with5.6*10^−12^ sterility levels when grown uder GH coditions. On the other hand P1 with 6.9*10^−11^ had lowest fertility levels compared to basmati370 that had 1.1*10^−4^ (highest) among the parents under OGH growth conditions. However, there was no significance difference in pollen sterility under GH and OGH growth conditions among all the parental lines (Table 1a). Some F1 seeds obtained in each cross breed are as recorded in Table 2

**Table 1:** Unpaired T-test analysis of % viable/fertile and sterility pollen. Table la shows pollen sterility under greenhouse environment (GH) while table lb shows pollen fertility under outside greenhouse (OGH) or natural growth conditions. Abbreviations P, T, P and B stand from PGMS, TGMS and basmati respectively.

| Varieties | GH growth condtions | *T*-test values | *p*-value (0.05) |
| --- | --- | --- | --- |
| P1 | Percentage pollen sterility | $2.4 \times 10^{-11}$ | 0.0001 |
| T | Percentage pollen sterility | $1.6 \times 10^{-10}$ | 0.0001 |
| P2 | Percentage pollen sterility | $3.3 \times 10^{-11}$ | 0.0001 |
| B217 | Percentage pollen sterility | $5.6 \times 10^{-12}$ | 0.0001 |
| B370 | Percentage pollen sterility | $3.9 \times 10^{-12}$ | 0.0001 |

b
| Varieties | OGH growth condtions | *T*-test values | *p*-value (0.05) |
| --- | --- | --- | --- |
| P1 | Percentage pollen fertility | $6.9 \times 10^{-11}$ | 0.0001 |
| T | Percentage pollen fertility | $5.7 \times 10^{-8}$ | 0.0001 |
| P2 | Percentage pollen fertility | $1.3 \times 10^{-8}$ | 0.0001 |
| B217 | Percentage pollen fertility | $2.5 \times 10^{-5}$ | 0.0001 |
| B370 | Percentage pollen fertility | $1.1 \times 10^{-4}$ | 0.0001 |

**Table 2:** Total number of F_1_ seeds produced.

| Line | *P1* | *P2* | T1 |
| --- | --- | --- | --- |
| B217 | 175 | 620 | 225 |
| B370 | 599 | 411 | 299 |

Over 80% of anthers locules from EGMS grown outside the greenhouse conditions, were filled with conspicuous pollen grains (Fig 2a), but locules for EGMS grown under greenhouse growth conditions had no observable pollen grains (Fig 2b). The EGMS grown OGH and inside GH had their staining yellow and spikelets had no observable grains (Figs 2a and b However, most pollen for EGMS OGH stained blue black and spites were conspicuous filled with grain (Figs 3 c and d).

**Fig. 3:**
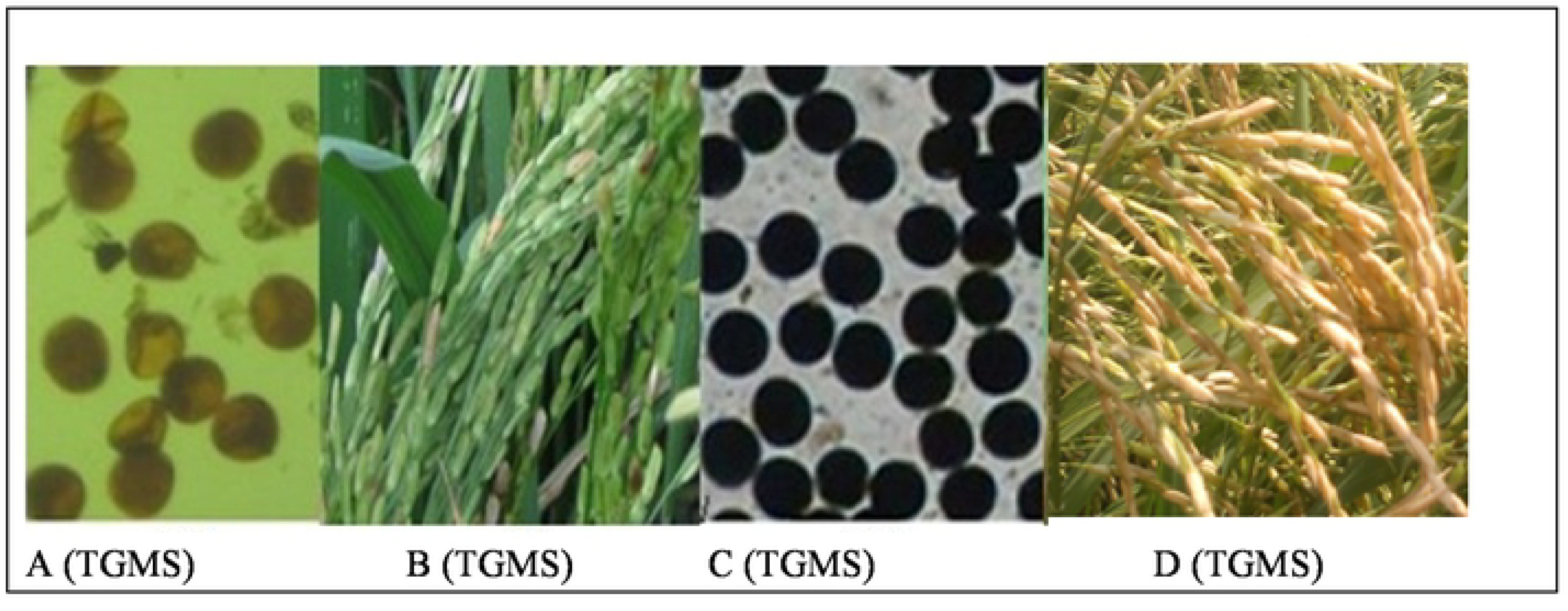
Comparison of pollen and seed set in GH and OGH condition (under X10 magnification). Figure (A) and (C) show pollen grains from GH and OGH while (B) and (D) show spikelets from plants grown under GH and OGH growth conditions respectively.

#### Evaluation of hybrid lines

Hybrids obtained from crosses between P1 x Basmati217, P1 x Basmati 370, T x Basmati217, T x Basmati370, P2 x Basmati217 and P2 x Basmati370 were coded P1B217, P1B370, TB217, TB370, P2B70 and P2B370 respectively. The hybrids were sown under GH environment where they were assessed for nine traits including number of productive tillers per hill, plant height (cm), days to 50% flowering, heading and maturity, panicle length and exertion (cm), percentage seed set and 1000 grain yield per plants (grams). Evaluation was based on standard evaluation system rice (25). Mean performance of parents and hybrids indicated high genetic variability in height (HT), maturity day (MD), 1000 seeds weight (1000SW), panicle exertion (PE), total spikelets (TS) and fertile spikelets (FS) (Tables 3 and 4).

**Table 3:**
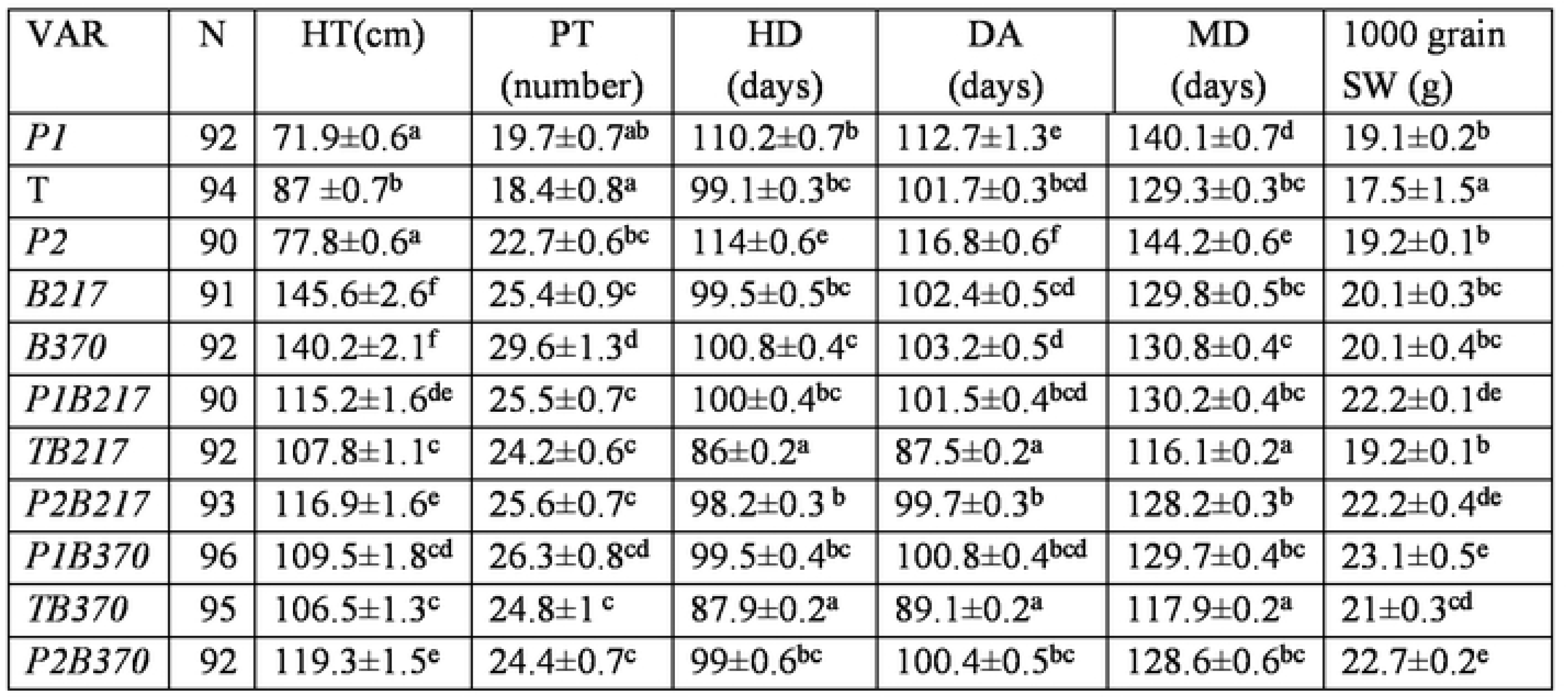
Evaluation of yield traits of hybrids lines. Values before ± sign are means of variables per plant. Means with different superscript letters within a column are significantly different (P < 0.05). N=number of plants sampled per variety. Variety =VAR, N= number of plants in each sample, Height = HT, Productive tillers =PT, Heading date=HD, Days to Anthesis =DA, Maturity date=MD and 1000 seed weight =SW.

**Table 4:** Hybrids and Parental varieties means of morphological traits. Values before ± sign are means of variables per plant. Means with different superscript letters within a column are significantly different (P < 0.05). SD = Standard deviation of the mean. Varieties (VAR), Panicle length (PL), Panicle exertion (PE), Total glumes (GL), Filled spikelets (FS), Sterile spikelets (SS), Percentage sterility (%S).

| VAR | N | PL(cm) | PE(cm) | GL | FS<br>(grains) | SS | %S |
| --- | --- | --- | --- | --- | --- | --- | --- |
| *P1* | 276 | 18.8±0.2<sup>a</sup> | 0.1±0<sup>a</sup> | 90±1.8<sup>b</sup> | 0.7±0.3<sup>a</sup> | 89.2±1.8<sup>ef</sup> | 99.3±0.3<sup>d</sup> |
| *T* | 282 | 23.5±0.2<sup>b</sup> | 0±0<sup>a</sup> | 116±1.9<sup>cd</sup> | 1.7±0.4<sup>a</sup> | 114.3±1.8<sup>g</sup> | 98.9±0.3<sup>d</sup> |
| *P2* | 270 | 19.2±0.1<sup>a</sup> | 0±0<sup>a</sup> | 96.3±1.5<sup>b</sup> | 0±0<sup>a</sup> | 96.3±1.6<sup>f</sup> | 100±0<sup>d</sup> |
| *B217* | 273 | 25.3±0.2<sup>c</sup> | 4.8±0.1<sup>c</sup> | 122.9±2<sup>cde</sup> | 37.1±1.4<sup>b</sup> | 85.8±2<sup>e</sup> | 69.5±1.1<sup>c</sup> |
| *B370* | 282 | 25.9±0.2<sup>cd</sup> | 3.8±0.1<sup>b</sup> | 126.9±1.7<sup>e</sup> | 34.3±1.3<sup>b</sup> | 92.7±1.9<sup>ef</sup> | 72.6±1.1<sup>c</sup> |
| *P1B217* | 270 | 25.3±0.1<sup>c</sup> | 5.9±0.1<sup>d</sup> | 128.2±1.5<sup>e</sup> | 68.5±1.2<sup>e</sup> | 59.7±1.2<sup>abc</sup> | 46.4±0.7<sup>a</sup> |
| *TB217* | 273 | 25.6±0.1<sup>c</sup> | 8.1±0.1<sup>f</sup> | 115.9±1.4<sup>c</sup> | 49.4±1<sup>d</sup> | 66.5±1.4<sup>cd</sup> | 57±0.8<sup>b</sup> |
| *P2B217* | 279 | 24.1±0.2<sup>b</sup> | 5.7±0.1<sup>d</sup> | 124.6±1.9<sup>de</sup> | 69.5±1.4<sup>e</sup> | 54.6±1.1<sup>a</sup> | 44.4±0.7<sup>a</sup> |
| *P1B370* | 279 | 25.2±0.2<sup>c</sup> | 6.8±0.3<sup>e</sup> | 126.4±2.1<sup>e</sup> | 53.6±2<sup>d</sup> | 72.8±1.8<sup>d</sup> | 58.8±1.1<sup>b</sup> |
| *TB370* | 285 | 26.5±0.1<sup>d</sup> | 6.7±0.1<sup>e</sup> | 85.8±1.7<sup>a</sup> | 42.9±1<sup>c</sup> | 63.6±1.4<sup>bc</sup> | 59.6±0.7<sup>b</sup> |
| *P2B370* | 282 | 24±0.1<sup>b</sup> | 5.6±0.1<sup>d</sup> | 114.3±2.1<sup>c</sup> | 67±1.7<sup>e</sup> | 58.9±1.1<sup>ab</sup> | 47.5±0.7<sup>a</sup> |

Yield and morphological traits are analysed in Tables 3 and 4. Height of the cultivars, (Table 3), revealed that line T (TGMS) had 87 ±0.7^b^cm, P1 (PGMS) had 71.9±0.6^a^cm and P2 had 77.8±0.6^a^cm. Basmati217 and B370 were the tallest plants with (145.6±2.6^f^; 140.2±2.1^f^) respectively. Hybrids height ranged between 106.5±1.3^c^ cm to 119.3±1.5^e^cm with P2 xB370 being tallest and T x B370 being the shortest (Table 4). The EGMS lines had lowest number of productive tillers (PT) while B370 had the highest (29.6±1.3^d^), and F_1_’s had an average of (24.2±0.6^c^ to 26.3±0.8^cd^) tillers (Table 3). Among the hybrids P1x B370 recorded the highest number of PTs. Lines PGMS (P1 and P2) had longest days to heading (HD), anthesis (AD), and maturity (MD), followed by TGMS (T) and Basmati had shortest (Table 3). Hybrids lines P1xB217, P1xB370, P2xB370, P2 xB217 and TxB370 had heavier seeds ranging between (21±0.3^cd^-23.1±0.5^e^) grams and also exceeding their respective parental lines. Line TxB217 had the lowest weight of 19.2±0.1^b^grames among hybrids. The 1000 grain weight of T, P1 and P2 lines was 17.5±1.5^a^, 19.1±0.2^b^ and 19.2±0.1^b^, respectively and those of pollen donor basmati370 and 217 were 20.1±0.3^bc^; 20.1±0.4^bc^ respectively (Table 4). Hybrid line TB370 had the longest panicles (PL) of 26.5±0.1^d^ followed by B370, 25.9±0.2, EGMS varieties had the shortest panicles, while other lines had almost similar values ranging from (24.1±0.2^b^to 25.3±0.2^c^) (Tables 3 and 4).

Lines P1, P2 and T had over 98% spikeletes sterility while basmati370 and 217, and F_1_’s cultivars had over 69% sterility (Table4). Hybrid lines P2B217 (56%), P1B217 (54%) and P2B370 (53%) (Table 4). had significantly lower sterility than the parents. Line P1B217 with (128.2±1.5^e^) had the highest number of spikelets counted. The EGMS had no measurable panicle exertion or uppermost internode. The F_1_’s had longest panicle exertions with uppermost internode measuring between (5.6±0.1^d^ to 8.1±0.1^f^) followed by the basmati370 and 217 that had an exertion of (4.8±0.1^c^) and (3.8±0.1^b^) respectively. On average TB370 had the lowest number of spikelets among the lines followed by P1 and P2. Spikelet length of basmati, T and hybrids (P1B217, P1B370, P2B370, P2B217, and TB217) ranged between (114.3±2.1^e^-128.2±1.5^e^ cm). Total glumes (GL) for B370, P1B21, P1B370 with 126.9±1.7e, 128.2±1.5e, 126.4±2.1e was significantly higher than the EGMS lines. Filled spikelets for three hybrids, P1B217, P2B217, and P2B370 with 68.5±1.2^e^, 69.5±1.4^e^ and 67±1.7^e^ were significantly higher than all parents. The three had the lowest sterility percentage (Table 4).

#### Correlating parental and hybrid phenotypic traits

Phenotypic correlations among F_1_’s namely P1B217, P1B370, P2B370, P2B217, TB217 and TB370 are shown in Table 5. Plant height (PH) corrected with productive tillers and with seed weight with values of r= 0.286 and r= 0.336 respectively. Heading days positively correlated to AD and MD with values of r= 0.986 and r= 0.967 respectively. Other positive correlations were observed between AD and MD (r=.967**), and PT and seed weight with r= 0.195.

## Discussion

Environment-sensitive genic male sterile (EGMS) rice, both PGMS and TGMS, grown under temperature higher than 34°C in the greenhouse had over 98% of their pollen staining blue-black with 1% potassium iodide (Figs 1 and 2). This is an indication that their pollen were completely male gamete sterile (26) and thus cannot have self-fertilization at this time. Therefore, EGMS can be pollinated with a pollen donor to produce hybrid seeds without adulteration from self-bred seeds. Under similar GH growth conditions basmati370 and 217 had over 20% of pollen staining blue-black, an indication that GH growth conditions could not induce complete male gamete sterility among them. The PGMS are male gamete sterile when grown under a long day of over 13.5 hours daylight length and high temperature can compensate for slightly shorter daylight length (26). In this study, PGMS were grown under GH growth and under 12hour-day length growth conditions and over 98% of pollen stained yellow in colour, an indication that they were sterile. Thus, high temperature compensated for the long day light length requirement for induction of complete pollen sterility in PGMS lines P1 and P2. Yuan (19) reported that, high temperature reduces the photoperiod required to induce complete male sterility in PGMS.

Greenhouse induced day-time temperature of above 34 °C was able to completely induce male sterility among P1, P2 and T with over 98% sterility (Tables 2-3). Many of the pollen were of abortive type and it stained yellow with 1% potassium iodide. EGMS exposed to high temperature had as low as less 2% seed set rate. It means use of staining method is accurate method of monitoring spikelet fertility. The EGMS exposed to temperature of around 24 °C under natural environment (OGH), at the time of critical sterility point, recorded some fertile pollen (Figs 3c and d). Within this temperature range EGMS revert to fertility, a time when they can propagate themselves (27, 18). For TGMS line T grown under high greenhouse (GH) temperature conditions, only 2% pollen fertility was realized. This was insignificant compared to PGMS lines P1 and P2. The results for basmati 370 and 217 grown under the GH growth conditions indicated that, they had significantly higher seed set rate than EGMS (Fig 1). This is an indication that they do not have thermo/photo sensitive male sterility genes like the EGMS. Therefore, they can be used as pollen donor in hybrid rice production programme.

Pollen sterility in lines P1, P2 and T grown under GH was over 97% and with a seed set rate of less 2% (Table 4). Thus, there was an inverse correlation between pollen sterility and seed set rate. This observation is affirmed by Ku, et al. (28) who reported that TGMS and PGMS lines grown under high temperature growth condition have significantly reduced pollen fertility at p>0.05. Lines P1 and P2 are PGMS while T is a TGMS. PGMS sterility responds to long photoperiod day length. Temperature of over 34°C completed induced both the PGMS and TGMS to complete sterility under light day length of 12hours. This is also an indication that high temperature can effectively compensated for long-day-light length requirement by PGMS lines to realize 100% sterility. Elevated temperatures can prevent adulteration of hybrid seeds with self-bred during cross-breeding (26).

Unpaired *t-*test results in both GH and OGH growth environments had a significance variance at p≤0.05for days to heading (Table 5). The EGMS varieties P1 had the highest p-value followed by P2 (Table 1). Also, sterility is influenced by the level of temperature which influences the overall level of pollen viability (Fig 1). This explains why lines P1, T, and P2 did not have seeds under GH growth conditions, unlike the ones grown under natural environment, and pollen donors lines basmat370 and 217 (Figs 2 and 3).

**Table 5:** Pearson correlation coefficients of plant height (PH), productive tillers (PT),heading day (HD), anthesis day (AD), maturity day (MD) and 1000 seed weight per plant (SW). Days to heading (HD), anthesis (AD) and maturity (MD) had high relationship in all varieties studied i.e. r=986 to r=967 range. Their positive values were much close to one comparing with other parameters studied. **. Values in parenthesis indicate correlation is significant at the P<0.01.

|  | PH | PT | HD | AD | MD | SW |
| --- | --- | --- | --- | --- | --- | --- |
| PH | 1 | .286** | -.296** | -.295** | -.298** | .336** |
| PT |  | 1 | -.109** | -.117** | -.109** | .195** |
| HD |  |  | 1 | .986** | .980** | -.126** |
| AD |  |  |  | 1 | .967** | -.167** |
| MD |  |  |  |  | 1 | -.125** |
| SW |  |  |  |  |  | 1 |

Lines TB217 and TB370 were better than the rest in anthesis (AD), days to heading (HD), and days to maturity (MD) (Table 3). Grain weight for P1B217, P2B217, and P1B370 was significantly higher than that of all parents. Line P1B370 had significantly higher productive tillers and panicle length than all other parents apart from B370. All hybrids had significantly larger panicle exertion than the parents. Good panicle exertion facilitates harvesting and cross pollination. Two hybrid lines P1B217 and P1B370 had a significantly higher total glumes than the parents, while P1B21, P2B217 and P2B370 had better grain filling than all parents. Also, P1B217, P2B217, TB370 and P2B370 recorded least sterility, better than all parents. In percentage sterility, all hybrids recorded superior performance than the best parent, a condition referred to as heterobeltiosis (29). In all other traits, hybrid were intermediately between the two parents apart from glume length in TB370, with a 85.8±1.7^a^, that was below the least performing parent.

All hybrids, apart from TB217weighed heavier than EGMS and Basmati parental lines. Increase of the grain weight also increases rice yield. Heavier seeds are preferred because they are healthy with more nutrients and when planted they result to vigorous seedlings with more roots, ability to with stand harsh conditions such as drought. On the other hand, small seeds are associated with reduced seedling vigour and also, difficult for mechanical harvesting. Grain length (GL), thickness and width determine grain size. The three traits; grain length (GL), thickness and width are quantitatively inherited and controlled by several genes (30). To date, it has been possible to isolate five key genes controlling seed size in rice namely: *GS3*, *GW2*, *qSW5* or *GW5*, *GIF1* and *GS5* (31–34). Gene *GS3* has a major effect on seed length, whereas *qSW5/GW5* and *GW2* confer both the seed or grain width (GW) and weight in rice. According to Yoshida, (35) and Sirajul (36) 1000 grain weight is a stable genetic character in rice.

Elongation of rice internodes is one of the most important traits for hybrid rice production which determines the plant height, pollination and underlies the grain yield (37). Panicle length (PL) and panicle exsertion (PE) exhibited variations under greenhouse condition with hybrids performing better than parental cultivars (Table5). Panicle length and panicle exertion in rice, are driven by uppermost internode elongation linked to internode elongation gene *eui1* (37). Complete panicle exsertion is under *eui1*gene and is influenced by temperature variations (38–40). Studies by Bardhan, et al. (41), Yang, et al. (38) suggest that different temperatures induce expression of male sterile gene in P(T)GMS lines at different levels. On the other side, the lower the temperature, the higher the expression level of *eui*gene, and the better the panicle exsertion, thus increased efficiency of cross breeding.

Some degree of F1 sterility was observed from crosses between EGMS and Basmati lines. Thus, yield can further be increased if this issue id addressed. This type of sterility has been observed in hybrid plants from *indicia* and *japonica* sub-species (16). According to Ikehashi and Araki, (5), certain *indica* and *japonica* hybrids show normal spikelet fertility in which case one or both parents possess a dominant wide-compatibility gene (*S5^n^*). Sterility and non-sterility is thought to be controlled by three alleles *S-5i* (in indica), *S-5j* (in japonica) and *S-5n* from WC rice (5, 16). According to Wan et al. (42), allelic interactions can be found at loci *S7, S8, S9, S15* and *S16* respectively, on chromosomes 4, 6, 7, 12 and 1. All of them cause sterility independent of each other (42). Genotypes *S-5n/S-5i* and *S-5n/S-5j* results in fertile female gametes but the *S-5i/S-5j* genotype produces semi-sterile panicles because of the partial abortion of female gametes, and this is what is postulated to have worked in this study as evidenced by the overall percentage seed set rate that was lower in hybrids than expected.

Basmati370 and 217 were taller than all the maternal parents but, EGMS and the hybrids displayed intermediate heights. These results are in line with the findings of Tua, et al. (43) and Kanya, et al. (44) who reported that hybrid rice had intermediary heights compared to their parents. Nevertheless, this is affected by cultivar type, agro-ecosystems involved and the cultural agronomic practices applied (45). Production of hybrids with intermediate heights in this study is significant in that, it can be utilized to breed for plants shorter than Basmati rice hence reduced lodging.

## Conclusion

High greenhouse temperatures of above 34°C during day time and 20°C at night can effectively emasculate both PGMS and TGMS varieties within Mwea Kenya (with 12hours of daylight length and 12hours of light length). This will allow production of basmati rice seeds in Kenya, using EGMS. Yield traits, such as grain weight showed better performance in hybrid than the best performing parent, thus, EGMS method can be used to increase yield in basmati370 and 217 through hybridization.

## Recommendations

The EGMS can be tested areas of Kenya hotter than Mwea to test ability to produce hybrids outside greenhouse growth conditions.

## Acknowledgement

National Commission of Science and Technology-Kenya (NCST) who funded the research.

## REFERENCES

1. FAO. Rice Market Monitor, FAO. 2017;

2. USAD. Foreign Agricultural service, Grain Report, Agricultural Information network, Grain and Feed Annual. Kenya Corn, Wheat and Rice Report. 2017.

3. Hinge VR, Patil HB, and Nada AB. Aroma volatile analyses and 2AP characterization at various developmental stages in Basmati and Non-Basmati scented rice (Oryza sativa L.) cultivars, Rice. 2016; 9:38: DOI 10.1186/s12284-016-0113-6

4. Emongor RA, Mureith FM, Ndirangu SN, Kitaka DM, Walela BM. The rice value chain in Kenya with reference to rice producers. 2009; KARI; Nairobi.

5. Kikuchi F. and Ikehashi H. Semidwarfing genes of high-yielding rice varieties in Japan. Rice Genetics Newsletter. 1984; 1: 93.

6. Peng, S, Tang, S Huang, J. Zou, Y. Cui, K. Zhang, Y. He, F. Laza, RC. and. Visperas RM Accelerating Hybrid rice development, F. Xie and B. Hardy (Ed): Yield attributes and nitrogen-use efficiency of “super” hybrid rice. IRRI. 2010; 420–428.

7. Khush GS. What it will take to Feed 5.0 Billion Rice consumers in 2030, Plant Molecular Biology. 2005; 59:1–6 DOI 10.1007/s11103-005-2159-5.

8. Kropff M, Cassman K, Peng S, Matthews R, Setter T. Quantitative understanding of yield potential.In: In: K. Cassman, Editor. Workshop on Rice Yield Potential in Favorable Environments Los Baños, Philippine; International Rice Research Institute; 1994. pp 21–38.

9. Yuan LP. The execution and theory of developing hybrid rice. Zhonggue Nongye Kexue (Chin. Agric. Sci.). 1977; 1:27–31.

10. Zhou G, Chen Y, Yao W, Zhang CJ, Xie WB, Hua JP, Xing JH, Xiao YZ and Zhang QF Genetic composition of yield heterosis in an elite rice hybrid. PNAS. 2012; 109 (39): 15847–15852.

11. Khush GS. Increasing the genetic yield potential of rice: prospects and approaches. International Rice commission Newsletter. 1994; 43: 1–7.

12. Wang Z, Han Q, Zi Q, Lv S, Qiu D, Zeng H Enhanced disease resistance and drought tolerance in transgenic rice plants overexpressing protein elicitors from Magnaporthe oryzae. PLoS ONE. 2017; 12(4): e0175734. https://doi.org/10.1371/journal.pone.0175734.

13. Huang M, Tang QY, Ao HJ, Zou YB. Yield potential and stability in super hybrid rice and its production Strategies. Journal of Integrative Agriculture. 2017; 16(5): 1009–1017.

14. Yuan LP. Progress in super-hybrid rice breeding, The Crop Journal. 2017; 5:100–102

15. Zhang Q, Liu KD, Yang GP. Molecular marker diversity and hybrid sterility in indica-japonica rice crosses. Theoretical Applied Genetics. 1997; 95: 112–118.

16. Yan L. China’s ‘super hybrid’ rice expected to yield 17 tons per hectare, People’s Daily Online, April 13, 2017;.

17. Virmani SS, Sun ZX, Mou TM, Jauhar AA, Mao CX. Hybrid rice and heterosis. Two-line hybrid rice breeding manual. Los Baños (Philippines): Breeding International Rice Research Institute. 2003;.

18. Shi MS and Deng JY. The discovery, determination and utilization of the hubei Photosensitive Genic Male-sterile Rice (*Oryzasativa*subsp. japonica). Acta Genetica Sinica. 1986; 13(2):107–112.

19. Yuan SC, Zhang ZG, He HH, Zen HL, Lu KY, Lian JH, Wang BX. Review and interpretation of two photoperiod-reactions in photoperiod-sensitive genic male-sterile rice. Crop Science. 1993; 33(4):651–660.

20. Ali J, Siddiq EA, Zaman FU, Abraham MJ, Ahmed I.. Identification and characterization of temperature sensitive genic male sterility sources in rice (Oryza sativa L.). Indian Journal of Genetics, 1995; 55 (3): 243–259.

21. Cao L, Zhan X. Chinese Experiences in Breeding Three-Line, Two-Line and Super Hybrid Rice. INTECH, 2014; Available at: http://creativecommons.org/licenses/by/3.0/PDF.

22. Dela Cruz N, Khush GS. Rice grain quality evaluation procedures. In: Singh RK, Singh US, Khush GS, editors. Aromatic rice. Oxford and IBH Publishing Co: Pvt. Ltd; 2000; New Delhi, India.

23. Njiruh PN and Xue Q. Tracking the expression of photosensitive genic male sterility genes in rice.African Journal of Biotechnology. 2013; 12 (47): 6583–6590.

24. Faize FA, Sabar M, Awan TH, Ijaz M, Manzoor Z. Heterosis and combining ability analysis in basmati rice hybrids. 2006; J. Anim. Pl. Sci. 16(1-2): 56–59

25. International rice research institute. Standard evaluation system for rice, 2002; pp 1–58

26. Njiruh PN and Xue Q..Programmed cell death-like behavior in photoperiodsensitive genic male sterile (PGMS) rice. African Journal of Biotechnology. 2011; 10(16):3027–3034.

27. Kumar P and Nautiyal MK. Two-Line Hybrid Production System and Their Applications in Rice. International Journal of Agriculture Sciences. 2016; 8(61):3502–3504.

28. Ku C, Kim B, Chung S.. Cytological observation of two environmental genic male sterile lines of rice. Molecular Cell. 2001; 12 (3):403–406.

29. Meena HS, Ram B, Kumar A, Singh BK, Meena PD, Singh VV and Singh D. Heterobeltiosis and standard heterosis for seed yield and important traits in Brassica juncea Journal of Oilseed Brassica. 2014; 5(2) 134–140.

30. Li Y, Fan C, Xing Y, Jiang Y, Luo L, Sun L, Shao D, Xu C, Li X, Xiao J, He Y, Zhang Q. Natural variation in GS5 plays an important role in regulating grain size and yield in rice. Nature Genetics. 2011; doi:10.1038/ng.977.

31. Fan C, Xing Y, Mao H, Lu T, Han B, Xu C, Li X, Zhang Q. GS3, a major QTL for grain length and weight and minor QTL for grain width and thickness in rice, encodes a putative transmembrane protein. Theoretical Application Genetics. 2006; 112: 1164–1171.

32. Song XJ, Huang W, Shi M, Zhu MZ, Lin HX. A QTL for rice grain width and weight encodes a previously unknown RING-type E3 ubiquitin ligase. Nature Genetics 39. 2007; 623–630.

33. Shomura A, Izawa T, Ebana K, Ebitani T, Kanegae H, Konishi S, Yano M. Deletion in a gene associated with grain size increased yields during rice domestication. Nature Genetics, 2008; 40: 1023–1028.

34. Weng JF, Gu SH, Wan XY, Gao H, Guo T, Su N, Lei CL, Zhang X, Cheng ZJ, Guo XP, Yang Z, Sun X, Wang S, Zhang Q. Genetic and physical mapping of a new gene for bacterial blight resistance in rice. Theoretical Applied Genetics, 2003; 106: 467–1472.

35. Yoshida S. Fundamentals of Rice Crop Science. Los Baños, Leguna, Philippines: International Rice Research Institute: 1981; pp 269.

36. Sirajul MI, Shaobing P, Romeo MV, Sultan MU, Hossain,SM, Julfiquar AW. Comparative study on yield and yield attributes of hybrid, inbred, and npt rice genotypes in a tropical irrigated ecosystem. Bangladesh Journal of Agricultural Research. 2010; 35(2): 343–353.

37. Huihai X, Youwei Y, Chunxia C, Jingzhen C. Elongation of the uppermost internode for Shuangdi Pei164s, TGMS rice with *eui*gene. African Journal of Agricultural Research. 2012; 26: 3806–3812.

38. Hittalmani S, Huang N, Courtois B, Venuprasad R, Shashidhar HE, Zhuang JY, Zheng KL, Liu GF, Wang GC, SidhuJS, et al. Identification of QTL for growth- and grain yield-related traits in rice across nine locations of Asia; Theoretical and Applied Genetics. 2003; 107:679–690.

39. Yang Z, Sun X, Wang S, Zhang Q. Genetic and physical mapping of a new gene for bacterial blight resistance in rice. Theoretical Applied Genetics, 2003; 106: 467–1472.

40. Ma A, Nawab NN, Abbas A, Zulkiffal M, Sajjad M. Evaluation of selection criteria in Cicerarietinum L. using correlation coefficients and path analysis. Australian Journal of Crop Science. 2009; 3: 65–70.

41. Bardhan RSK, Pateña GF, Vergara BS. Feasibility of selection for traits associated with cold tolerance in rice under rapid generation advance method. Euphytica, 1982; 31: 25–31.

42. Wan J, Yamaguchi Y, Kato H, Ikehashi H. Two new loci for hybrid sterility in cultivated rice (Oryza sativa L.). 1996; Theoretical and Applied Genetics, 92: 183–190.

43. Tua S, Luana L, Liua Y, Longa W, Konga F, Hea T. Production and heterosis analysis of rice autotetraploid hybrids. 2007; Crop Science,47, 2356–2363.

44. Kanya JI, Njiruh NP, Kimani JM, Wajogu RK, Kariuki SN Evaluation of photoperiod and thermosensitive genic male sterile lines for hybrid rice seeds production in Kenya. International Journal of Agronomy and Agricultural Research (IJAAR). (2013) 3 (2):21–39. SSN 22237054 Dio-22253610.

45. Zafar N, Aziz S, Masod S. Phenotypic divergence for agro-morphological traits among landrace genotypes of rice *(Oryzasativa L.)* from Pakistan.International Journal of Agriculture and Biology. 2004; 2: 335–339.

